# Harnessing micrometer-scale tPA beads for high plasmin generation and accelerated fibrinolysis

**DOI:** 10.1101/2024.11.06.621942

**Authors:** Matthew J. Osmond, Fabrice Dabertrand, Nidia Quillinan, Enming J. Su, Daniel A. Lawrence, David W.M. Marr, Keith B. Neeves

## Abstract

Rapid restoration of blood flow is critical in treating acute ischemic stroke. Current fibrinolytic therapies using tissue plasminogen activator (tPA) are limited by low recanalization rates and risks of off-target bleeding. Here, we present a strategy using tPA immobilized on micrometer-scale beads to enhance local plasmin generation. We synthesized tPA-functionalized beads of varying sizes (0.1 μm and 1.0 μm) and evaluated their efficacy. *In vitro* assays demonstrated that 1.0 μm tPA-beads generated higher plasmin generation compared to free tPA and 0.1 μm beads, overcoming antiplasmin inhibition and promoting a self-propagating wave of fibrinolysis. In a murine model of acute ischemic stroke, intravenous administration of 1.0 μm tPA-beads at doses nearly two orders of magnitude lower than the standard free tPA dose led to rapid and near-complete thrombus removal within minutes. This approach addresses kinetic and transport limitations of current therapies and may reduce the risk of hemorrhagic complications.

**Teaser:** Micrometer-scale tPA beads improve stroke treatment by enhancing local clot-dissolving action.

## Introduction

Reestablishing blood flow as quickly as possible is the goal of most thrombotic disorder treatment approaches. Among current strategies is fibrinolysis, the biochemical degradation of the fibrin fibers that provide the mechanical integrity of occlusive thrombi. This is accomplished with recombinant tissue plasminogen activators (tPA), with alteplase and tenecteplase being the only currently approved FDA drugs. When patients present within 4.5 hr of the onset of symptoms, intravenous (IV) tPA is the first-line treatment for acute ischemic stroke, often used in combination with mechanical thrombectomy where available, resulting in better long term functional outcomes and reduction in mortality.(*1, 2*) However, since the introduction of recombinant tPA’s more than 30 years ago, outcomes in ischemic stroke have not changed with only 10-20% of large vessel occlusions recanalized with IV tPA,(*3*) highlighting the necessity of new approaches.

Plasminogen activators, and tPA in particular, are and have been the strategy of choice for fibrinolysis. tPA has a high affinity (K_d_=1-2 nM) to bind to fibrin fibers, thus it is a targeted therapy as it accumulates at the thrombus interface enhancing local plasminogen activation.(*4*–*6*) tPA’s endogenous inhibitor, PAI-1, circulates at a relatively low concentration (0.1-1 nM) and thus can be overcome by exogenous recombinant tPA with circulating concentrations of 10-50 nM following IV injection.(*7, 8*) Intracerebral hemorrhage and off target bleeding are the most common complications with tPA therapy, occurring in 3-8% of people receiving IV tPA for acute ischemic stroke.(*9, 10*) In addition, systemic activation of plasminogen by tPA promotes fibrinogenolysis and activation of enzymes and inflammatory pathways that can cause tissue damage and impede wound repair.(*11*–*15*)

The effectiveness of tPA is hindered by kinetic and transport challenges within thrombi. tPA is most effective when bound to fibrin, which serves as a co-factor to accelerate the activation of plasminogen to plasmin by 400-fold.(*6, 16, 17*) However, tPA and plasmin(ogen) compete for the same binding sites on fibrin fibers. This results in the counterintuitive observation that high tPA concentrations attenuate fibrinolysis, due to both competition for binding sites and plasminogen depletion. Indeed, plasminogen availability is rate limiting under high tPA concentrations.(*18*–*20*) Furthermore, the lysis pattern of tPA mediated thrombolysis tends to be anisotropic and incomplete; regions of high permeability within the center of the thrombus tend to lyse first leaving small channels that partially reestablish blood flow but conversely lead to slow surface erosion of the remaining wall bound thrombus.(*21, 22*)

Direct infusion of plasmin, which would bypass PA’s, is hampered by binding with one of its endogenous inhibitors, α2-antiplasmin, which circulates at a high plasma concentration (∼1 μM).(*23, 24*) Local delivery of plasmin has been attempted via catheter, with success in animal models and phase I clinical trials but has not been shown superior to tPA.(*25*) Immobilization of tPA to particles, which generates plasmin locally while sidestepping the competition between tPA and plasmin for fibrin binding sites, is an alternative. This approach has been implemented primarily on sub-micrometer scale particles, often decorated with fibrin or platelet targeting moieties or embedded with iron oxide to allow for enhanced accumulation or local hyperthermia using external magnetic fields.(*9, 26*–*31*) These tPA-nanoparticle constructs can increase reperfusion times in animal models of thrombosis faster than free tPA.(*32*)

Here, we introduce an innovative yet straightforward approach: immobilizing tPA on micrometer-scale beads to significantly enhance local plasmin generation. Unlike previous nanoparticle-based strategies, our method leverages the size-dependent plasmin generation of larger beads to overcome kinetic barriers and antiplasmin inhibition, leading to rapid and efficient thrombus dissolution both *in vitro* and *in vivo*. In a murine model of acute ischemic stroke, we show that these particles entrain themselves within thrombi, lysing them from the inside-out resulting in near complete removal of thrombotic material in minutes. This is accomplished by using tPA doses that are almost two orders-of-magnitude lower than what is needed for similar recanalization times with free tPA.

## Results

### Fibrinolysis with tPA-beads is size dependent

To provide a faithful comparison between free and immobilized tPA on beads, we use tPA activity (mU/mL) as measured by a tPA specific chromogenic substrate. A standard curve was generated for tPA activity (0-2 μg/mL) and used to titrate tPA-bead concentrations to obtain an equivalent activity per milligram of beads (Figure S1). The 0.1 μm tPA-beads had approximately half the activity per mg than those of the 1.0 μm tPA-beads. For all subsequent *in vitro* experiments, we use a tPA activity of 2.25 mU/mL for both free tPA and tPA-beads, which corresponds to 1 μg/mL (14 nM) free tPA, a concentration that is comparable to the circulating concentrations of alteplase in humans following initial bolus.(*33*)

For equivalent tPA activities (2.25 mU/mL), 0.1 μm tPA-beads showed similar fibrinolysis rates in plasma derived fibrin gels as free tPA as measured by turbidity (Figure 1A) with a lysis phase that began at ∼60 min and continued to end of the experiment (180 min) while not quite reaching full lysis. Note that the 0.1 μm tPA-bead does not reach the same maximum absorbance suggesting some lysis occurred before the fibrin gel could fully form. For 1 μm tPA-beads the lysis was notably faster; there was a rapid lysis phase from 11 min to complete lysis at 38 min on average.

**Figure 1.**
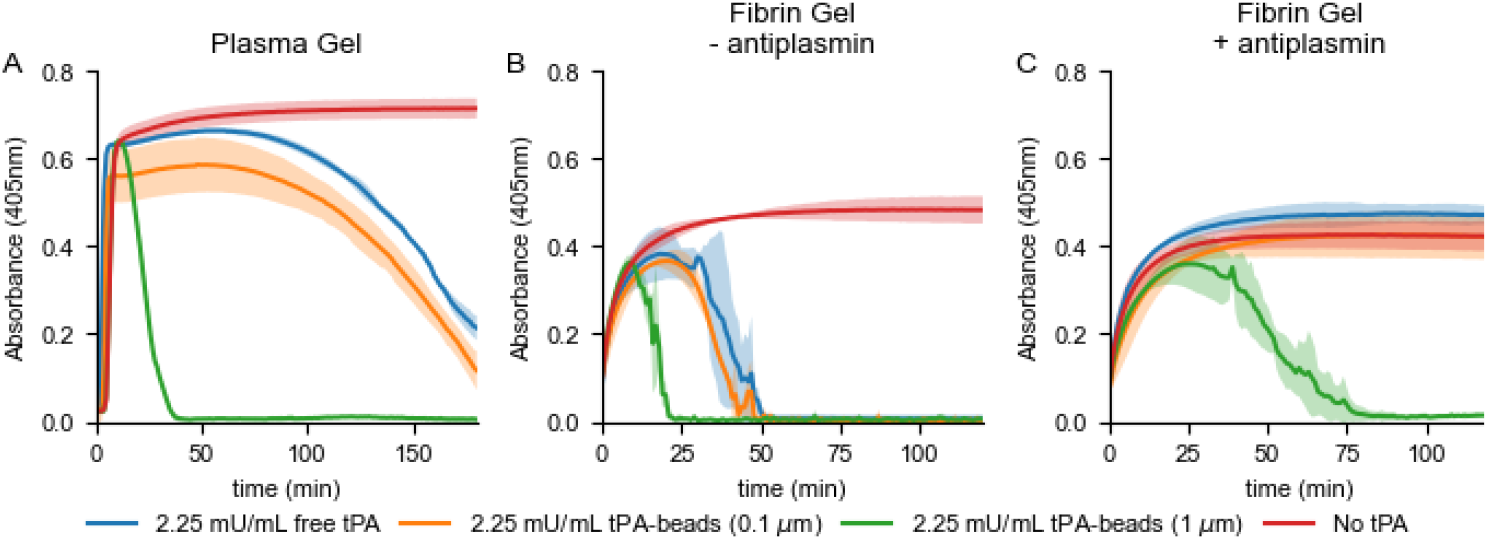
Size dependent fibrinolysis with tPA-beads. Fibrin gels were formed with (**A**) normal pooled plasma, tissue factor (1.25 pM) and CaCl_2_ (8.3 mM), (**B**) fibrinogen (2 mg/mL), glu-plasminogen (10 μM), thrombin (2 U/mL), (**C**) fibrinogen (2 mg/mL), glu-plasminogen (10 μM), thrombin (2 U/mL), α2-antiplasmin (1 μM) with free tPA or tPA-beads at 37°C. Fibrinolysis was measured by absorbance (405 nm) for tPA-beads (2.25 mU/mL) with diameters of 0.1 μm (3.55×10^10^ beads/mL) and 1.0 μm (1.69×10^8^ beads/mL) or free tPA (2.25 mU/mL). Each line and shaded region represent the average and standard deviation of n=3.

To determine if the differences between the 0.1 μm and 1 μm tPA-beads was a function of plasmin inhibition by antiplasmin, the same experiment was repeated in a purified system wherein a fibrin gel was formed from fibrinogen, thrombin, and plasminogen with or without antiplasmin at its plasma concentration (1 μM) (Figure 1B, C). In the absence of antiplasmin, the trends are similar to the plasma gel; the 1 μm tPA-beads have a shorter lag time and faster lysis rate compared to 0.1 μm tPA-beads and free tPA. In the presence of antiplasmin, only the 1 μm tPA-beads can induce fibrinolysis albeit at a prolonged lag time and rate compared to experiments without antiplasmin. These data suggest that plasmin generation on 1 μm tPA-beads can bind to fibrin fibers and overcome inhibition by antiplasmin more readily than 0.1 μm tPA-beads.

### Local plasmin flux from tPA-bead surface dictates global fibrinolysis

To determine how plasmin generation on bead surfaces compared to tPA activity, cleavage of a plasmin specific chromogenic substrate was measured. Here, a suspension containing 2.25 mU/mL tPA activity of 0.1 and 1 μm tPA-beads was compared to free 2.25 mU/mL free tPA (Figure 2). In the absence of any co-factors, the plasmin generation, as calculated by the slope of the kinetic absorbance curve, on 1 μm tPA-beads was about two-fold higher than 0.1 μm tPA-beads and free tPA with equivalent activity. In the presence of co-factors of tPA, fibrinogen and FDP, plasmin generation was significantly increased for all conditions. This observation suggests that even when the tPA is immobilized on the bead, it is still able to form a complex with these co-factors and plasminogen to increase the catalytic rate of the enzymatic reaction. However, the increased plasmin generation with these co-factors on 0.1 μm tPA-beads was not as prominent as for free and 1 μm tPA beads.

**Figure 2.**
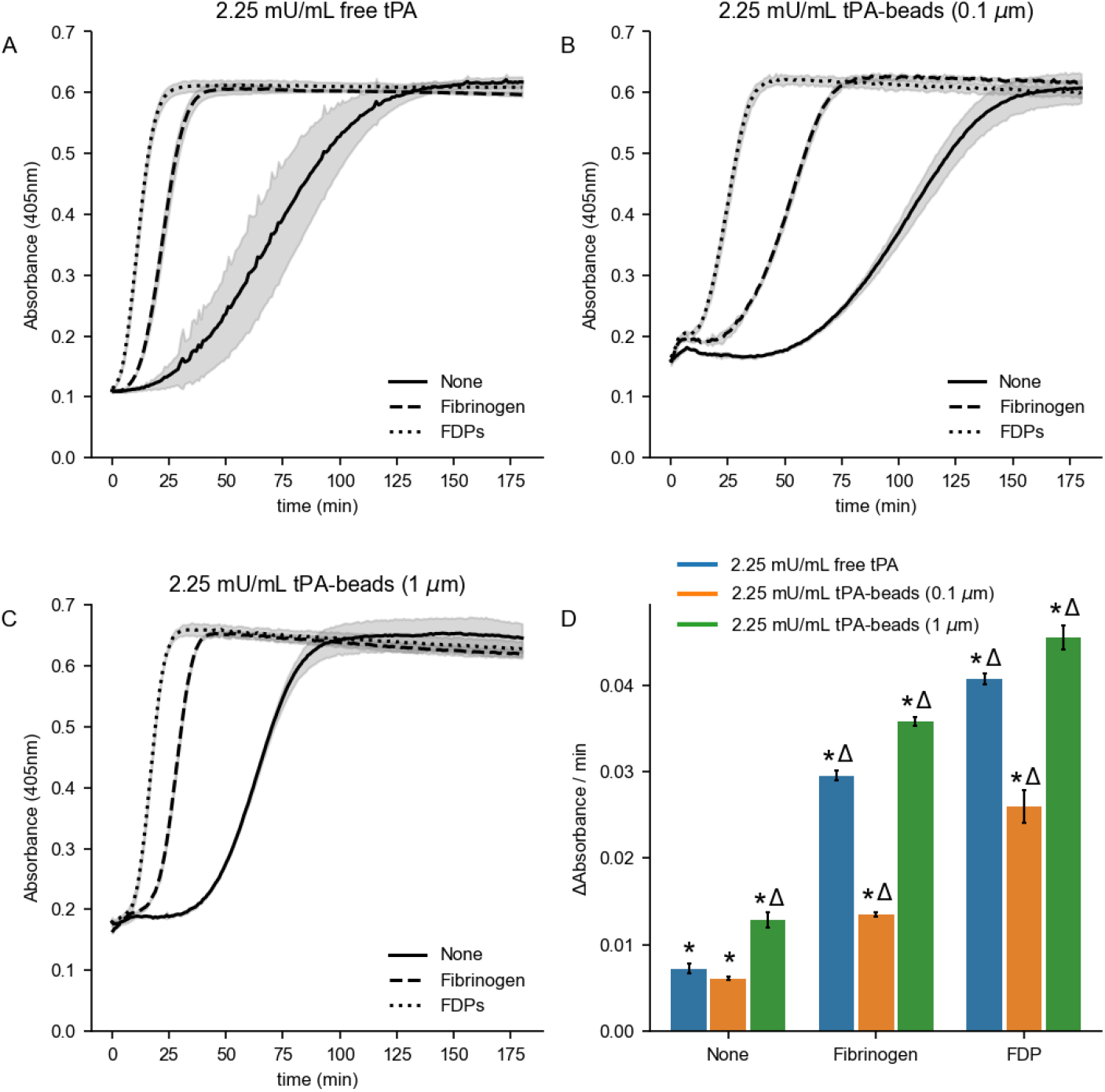
Plasmin generation of tPA-beads. Plasmin generation was measured using chromogenic substrate S-2251, glu-plasminogen, and 2.25 mU/mL tPA activity with (**A**) free tPA, (**B**) 0.1 μm t-PA beads, or (**C**) 1 μm tPA beads and its co-factors fibrinogen or fibrin degradation product (FDP) at 37°C. Each line and shaded region represent the average and standard deviation of n=3. (**D**) Slope of the reaction curves at 50% of the maximum absorbance for each condition. Bars and error bars represent the average and standard deviation of n=3. *, p < 0.001 represents significant difference between matching tPA conditions for a given co-factor (comparison of grouped blue, orange, and green bars). Δ, p < 0.001 represents the significant difference between co-factors for a given tPA condition (comparison of like colored bars).

### Micrometer tPA-beads catalyze self-propagating internal fibrinolysis

To explore fibrinolysis and the length scale of the tPA-beads, we observed fibrin fiber lysis around individual beads by confocal microscopy, using fluorescently labeled fibrin(ogen) and plasmin(ogen) (Figure 3). A more dilute tPA activity of 5.62 μU/mL of the 0.1 μm, and 1 μm tPA-beads was used here such that there was only a one bead per field of view. For both conditions, plasmin(ogen) appears to template on fibrin fibers and spreads radially from the bead. For the 0.1 μm tPA-beads, the templating occurs, but little lysis is observed over 1 hr.

**Figure 3.**
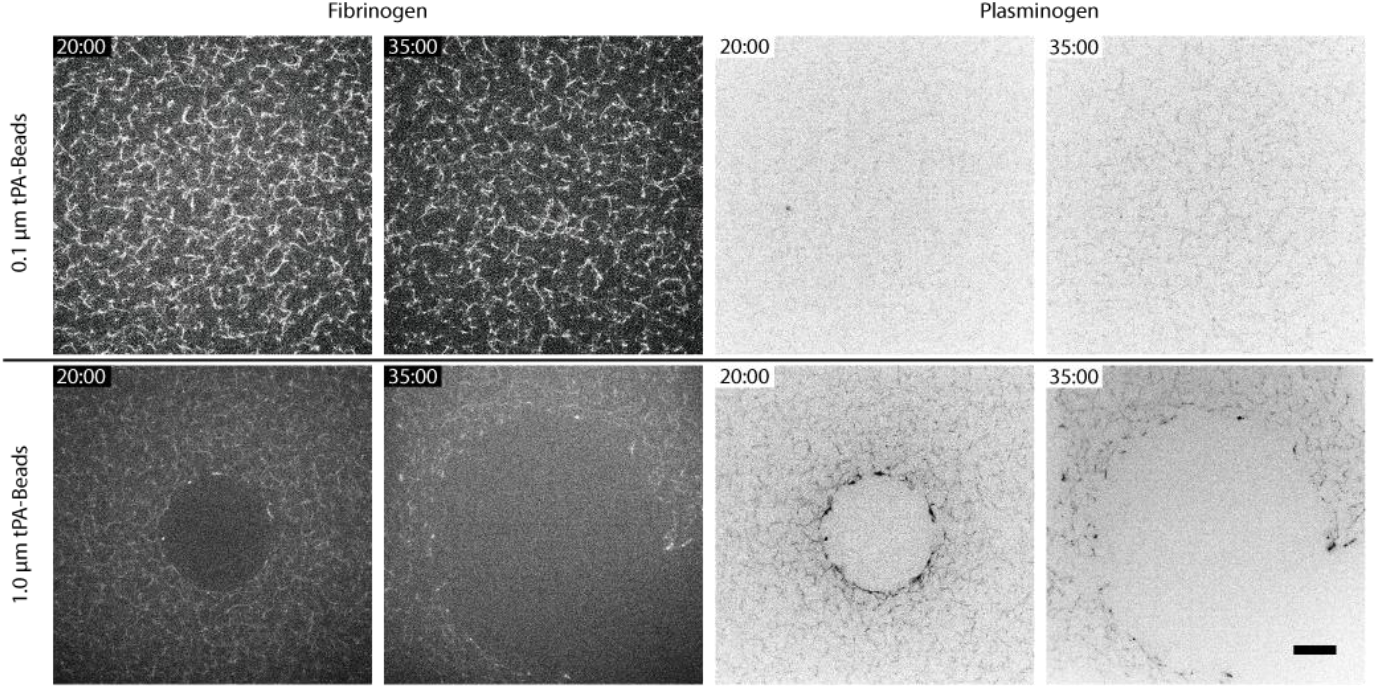
Internal fibrinolysis at the bead scale. Fluorescent micrographs of internal lysis of fibrin gels in a purified system (fibrinogen, thrombin, plasminogen) by 0.1 μm tPA-beads, or 1 μm tPA beads at 20 °C. The fibrin network was labeled using 40 μg/mL Alexa Fluor 647 labeled fibrinogen (1:25 labeled:unlabeled) and Pacific blue labeled glu-plasminogen (1:10 labeled:unlabeled). Time stamp in min:seconds. Scale bar = 20μm.

For the 1 μm tPA-beads, a halo of lysed fibrin forms around the bead, eventually propagating beyond the field of view and coalescing with halos of other nearby beads (Movie S1). The perimeter of the halo is enriched in plasmin(ogen) compared to the bulk solution. These halos, once initiated, self-propagate over a length scale of 100s of microns, much larger than the 1 μm beads. The diameter of these halos grows linearly with time suggesting this is not a diffusion-limited process which scales with square root of time (Figure 4). That is, plasmin does not appear to be diffusing from the tPA-bead to the lysis front but rather binding and unbinding to or crawling along fibrin fibers.

**Figure 4.**
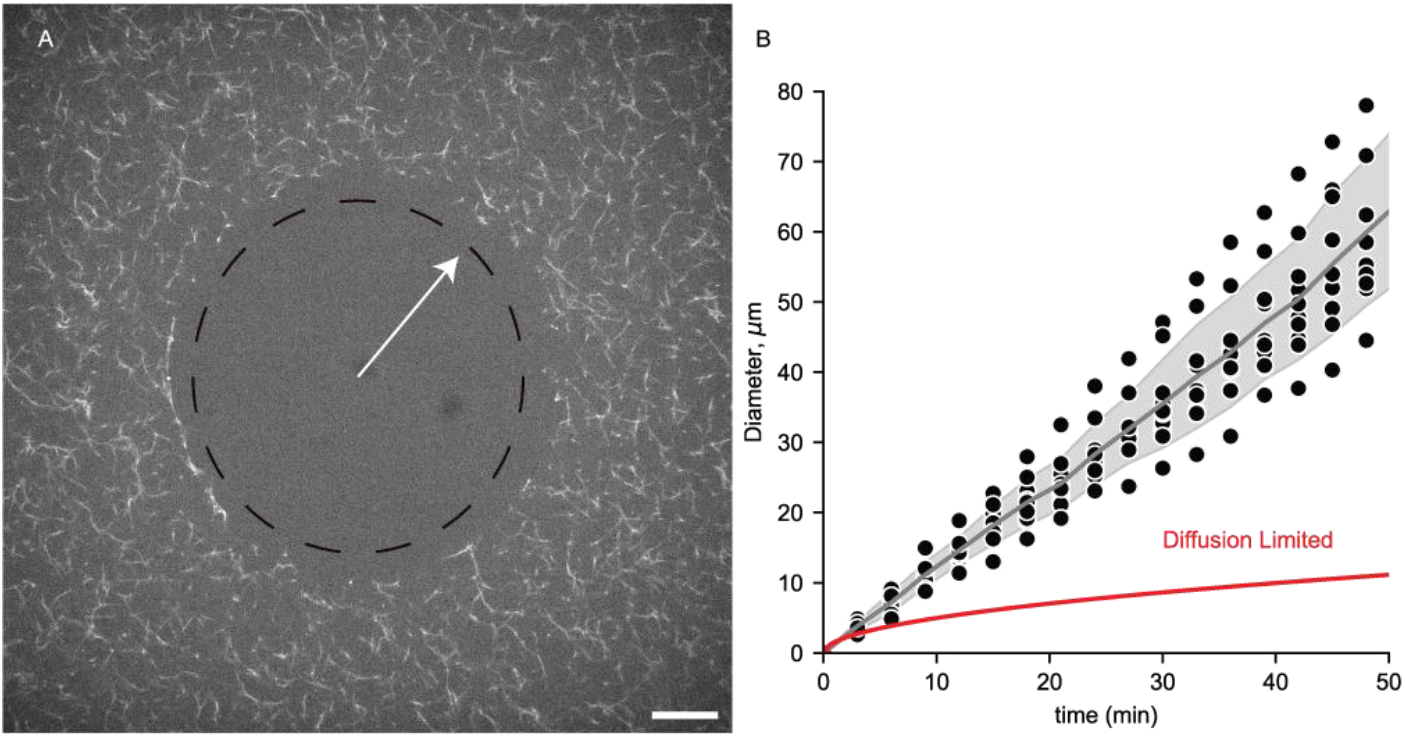
Growth rate of fibrinolytic halo around tPA-beads. (**A**) Fluorescent micrograph illustrating the lysis front around a 1.0 μm tPA-bead at t = 50 min. The fibrin network was labeled using 40 μg/mL Alexa Fluor 647 labeled fibrinogen (1:25 labeled:unlabeled) (**B**) Halo diameter as a function of time recorded from n =10 different tPA-beads. The line and shaded region represent the average and standard deviation. The red line represents a simple diffusion model with a diffusion coefficient that gives the same initial change in diameter as the real data.

### Simulation of plasmin flux and penetration

To aid in the interpretation of the single bead internal fibrinolysis assays (Figure 4), the plasmin flux from the surface of 0.1 and 1 μm tPA-beads and its penetration distance was estimated with a reaction-diffusion model. (Figure 5) In the model, plasminogen diffuses to the surface of the beads, is activated by tPA to plasmin, and plasmin diffuses away from the bead and, in the cases noted, binds to antiplasmin. The model solution is governed by two Dahmköhler numbers, which is a dimensionless ratio of the reaction rate to the diffusion rate (see Supplemental Materials). The first Dahmköhler number, Da_bead_, is the ratio of the reaction rate of plasminogen activation to plasmin by tPA to the diffusion rate of plasminogen to the bead surface. The second Dahmköhler number, Da_ap_, is the ratio of antiplasmin binding to plasmin to the diffusion rate of plasmin away from the bead surface. When the Da < 1, the process is reaction-limited, that is the reaction is the rate limiting step. Conversely, when the Da > 1, the process is diffusion-limited, and the diffusive transport is rate limiting.

**Figure 5.**
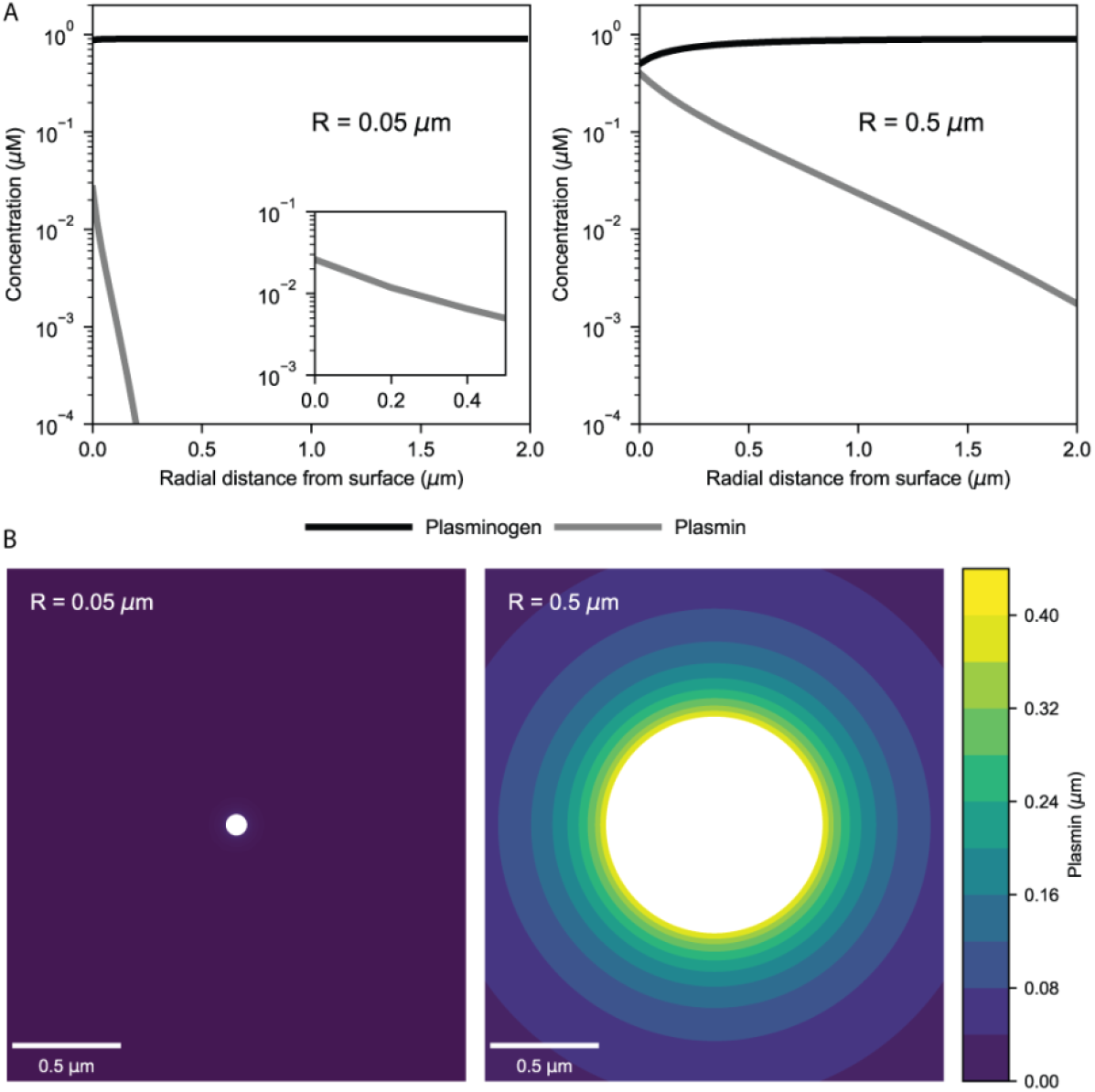
Simulation of plasmin generation on beads. A reaction-diffusion model of plasmin generation on tPA-beads is used to predict the concentration profiles (**A**) of plasminogen and plasmin as a function of radial distance from the bead surface in the absence of anti-plasmin. Contour plots (**B**) of plasmin concentration on 0.1 and 1.0 μm tPA-beads in the absence of anti-plasmin. Reaction constants used were calculated for a temperature of 20°C. Initial concentration of plasminogen was 0.9 μM.

For the conditions of the single bead lysis experiments (Figure 3), there is no antiplasmin, and the Da_bead_ = 0.20 for the 0.1 μm tPA-bead and Da_bead_ = 2.0 for the 1 μm tPA-bead. This suggests that there is transition in the dynamics of plasmin generation for the smaller bead, which is reaction-limited, to the larger bead, which is diffusion-limited. The factor of ten difference follows the linear scaling of Da_bead_ with bead radius. This transition from reaction-limited to diffusion-limited is reflected in the plasminogen concentration profile from each of two sized bead surface; there is no appreciable depletion of plasminogen near the 0.1 μm tPA-bead, while the near-surface plasminogen concentration is roughly half that of the bulk concentration. As the 1 μm tPA-bead creates a nearly two order of magnitude higher plasmin generation than 0.1 μm tPA-beads (260 fmol/s vs 4.6 fmol/s), the penetration of plasmin is significantly higher. For example, using 5 nM as a benchmark as this plasmin concentration lyses fibrin fibers in 10’s of min (*34*) similar to the time scale of our single bead lysis experiments, the model predicts this plasmin concentration 85 nm from the 0.1 μm bead and 1.9 μm radially from the 1 μm bead. Given that the pore size of fibrin or plasma gel made from 2-3 mg/mL of fibrinogen is on the order of 0.5-1.2 μm,(*35, 36*) these calculations suggest that plasmin generated on 0.1 μm tPA-beads are unlikely to reach fibrin fibers beyond those it is directly in contact with or within ∼100 nm. Conversely, 1 μm tPA-beads yield nanomolar concentrations at micrometer penetration depths likely affecting many nearby fibrin fibers.

To extend this analysis to the turbidity assays (Figure 1B, C), we add antiplasmin into the model. In this, the Da_bead_ values are within a factor of two due to temperature differences with the single bead lysis experiments (see Supplemental Materials); however, Da_ap_ is dramatically different for the two bead sizes. Da_ap_ = 0.43 for the 0.1 μm tPA beads and Da_ap_ = 43 for the 1 μm beads. This hundred-fold difference follows from the radius squared dependence of Da_ap_ and suggests that antiplasmin has little effect on plasmin penetration from smaller beads but dictates how far plasmin can penetrate from larger ones (Figure S2). This supports turbidity results showing an attenuation in lysis rate with 1 μm tPA-beads in the presence of antiplasmin. However, the model does not explain why 0.1 μm tPA-beads show no observable lysis.

### Micrometer tPA-beads accelerate external fibrinolysis compared to free tPA

The internal fibrinolysis experiments presented in the previous section show that when beads are mixed throughout a fibrin gel that 1 μm beads have faster lysis rate than the same activity of 0.1 μm tPA-beads and free tPA. To determine how tPA-beads lyse a fibrin gel when introduced at an interface, we used a ball sedimentation assay wherein the velocity of ball is used a proxy for fibrinolysis rate (Figure 6).(*37, 38*) In this, a 4.5 mU tPA bolus in 100 μL was placed on top of a fibrin gel for each condition (free tPA, 0.1 and 1 μm tPA-beads). This bolus was chosen to match that used in the bolus used in the photothrombotic murine stroke model (next section). As apparent in Fig. 6A, the 1 μm tPA-beads have a faster ball drop rate reaching the bottom of the cuvette within 200 min compared to the free tPA which takes 350 min (Fig. 6B). The 0.1 μm tPA-beads do not significantly lyse the clot over the observation period as the distance traveled is comparable to the saline negative control reflecting settling of the dense steel ball into the porous fibrin gel rather than any lysis.

**Figure 6.**
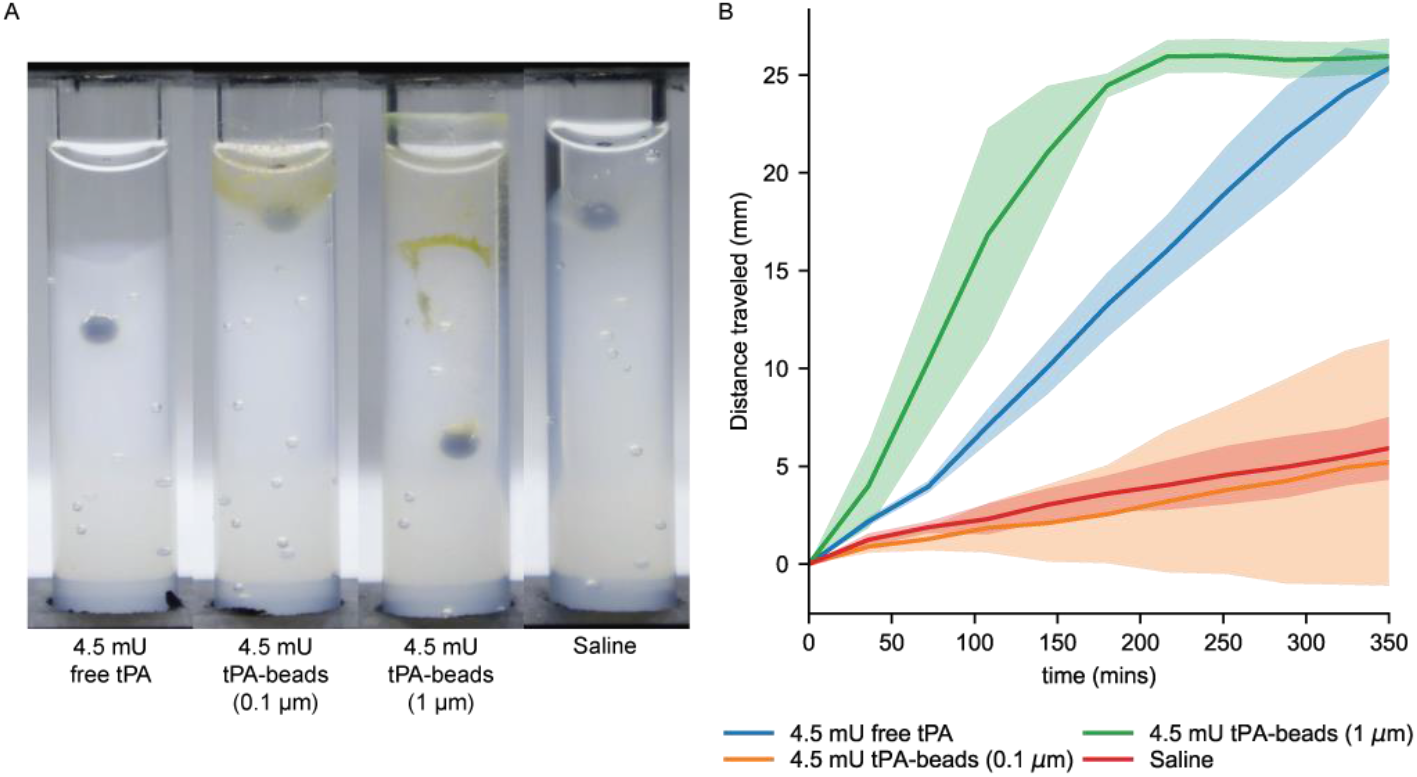
External fibrinolysis measured by the ball sedimentation assay. (**A**) Fibrin gels were formed using fibrinogen (2 mg/mL), plasminogen (0.2 μM), and thrombin (2 U/mL), then addition of a 100 μL bolus of (from left to right), 0.1 μm beads, 1 μm beads, and free tPA (each containing 4.5 mU tPA activity) and a saline only control. The photograph shows displacement of a 2 mm steel ball bead after 2 hr of lysis at 37°C. (**B**) Distance traveled for stainless steel beads over time. Each line and shaded region represent the average and standard deviation of n=3.

### Micrometer tPA-beads rapidly reestablish pre-occlusion blood flow in a photothrombotic stroke model

The *in vitro* experiments presented above show that 1 μm tPA-beads induce faster internal and external fibrinolysis than comparable concentrations or doses of free tPA and 0.1 μm tPA-beads and that they display a unique localized halo-like lysis pattern. To determine if these observations translate to *in vivo* thrombolysis, we used a photothrombotic thrombosis model in the MCA of the mouse. Saline, free tPA, tPA-beads (0.1, 1 μm), and beads (nonfunctionalized, 0.1, 1 μm) were delivered as a bolus through the tail vein. A tPA dose of 4.5 mU was chosen to reflect a systemic concentration comparable to *in vitro* experiments (2.25 mU/mL x ∼2 mL blood volume = 4.5 mU). We also included a 350 mU tPA dose (5 mg/kg) as this has been shown an effective dose for thrombolysis in this model.(*39*) Blood flow was measured by LDF and two metrics of thrombolysis were calculated, time to reperfusion and % pre-occlusion cerebral blood flow (Figure 7). Only the 4.5 mU dose of the 1 μm tPA-beads and the 350 mU dose of the free tPA showed measurable blood flow in the occluded vessel (Figure S3, Figure 7 B,C). In four out of the five trials with 1 μm tPA-beads, the time to reperfusion was ∼5 min, with one trial showing no reperfusion. In three of the five trials with 350 mU free tPA, the time to reperfusion was more variable and ranged from 5-15 min, with two trials showing no reperfusion. In the cases where reperfusion occurred for the 1 μm tPA-beads there was nearly complete rescue to blood flow to pre-occlusion levels. In contrast, for the 350 mU free tPA cases where there was reperfusion, there was 50-60% rescues of pre-occlusion blood flow.

**Figure 7.**
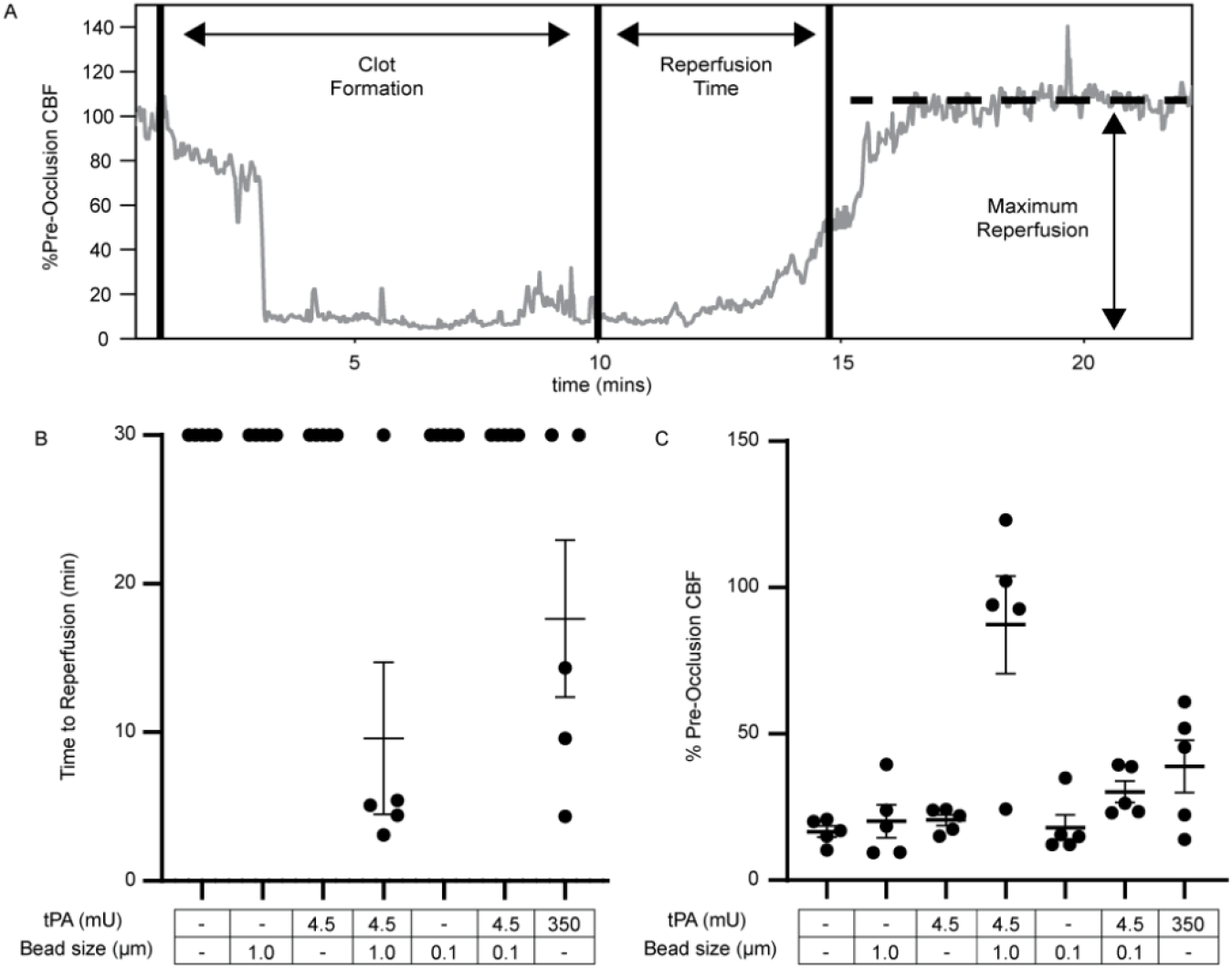
tPA-bead mediated thrombolysis in the murine photothrombotic stroke model. (**A**) A representative laser doppler flowmeter curve generated by measuring blood flow through the middle cerebral artery of a mouse during thrombus formation and lysis for a 4.5 mU dose of 1 μm tPA-beads. The time to reperfusion is defined as the time from the onset of the treatment to the inflection point where significant reperfusion occurs. The maximum perfusion is defined as the maximum achieved blood flow that happens after treatment onset. The (**B**) time to reperfusion and (**C**) maximum reperfusion for saline, 1.0 μm beads (no tPA), 4.5 mU free tPA, 4.5 mU 1.0 μm tPA-beads, 0.1 μm beads (no tPA), 4.5 mU 0.1 μm tPA-beads, and 350 mU free tPA.

To further explore these two conditions, intravital microscopy was used to visualize the thrombus and its dissolution (Figure 8). The lysis pattern with a 350 mU dose of free tPA was formation of small canulae that eventually re-canulated the vessel but, upon restoration of blood flow, showed little additional lysis of the thrombus attached to the vessel wall over the observation period of 30 min (Movie S1). The 4.5 mU dose of 1 μm tPA-beads caused the entire thrombus to slough off the vessel wall and be completely removed by blood flow (Movie S2). Following removal of the occlusive thrombus subsequent blood cell accumulation on the vessel wall that was also removed within minutes. Fluorescently-labeled 1 μm beads without tPA were injected and found to entrain themselves throughout the thrombus, suggesting that tPA-beads were able to induce lysis from the inside out (Figure S4).

**Figure 8.**
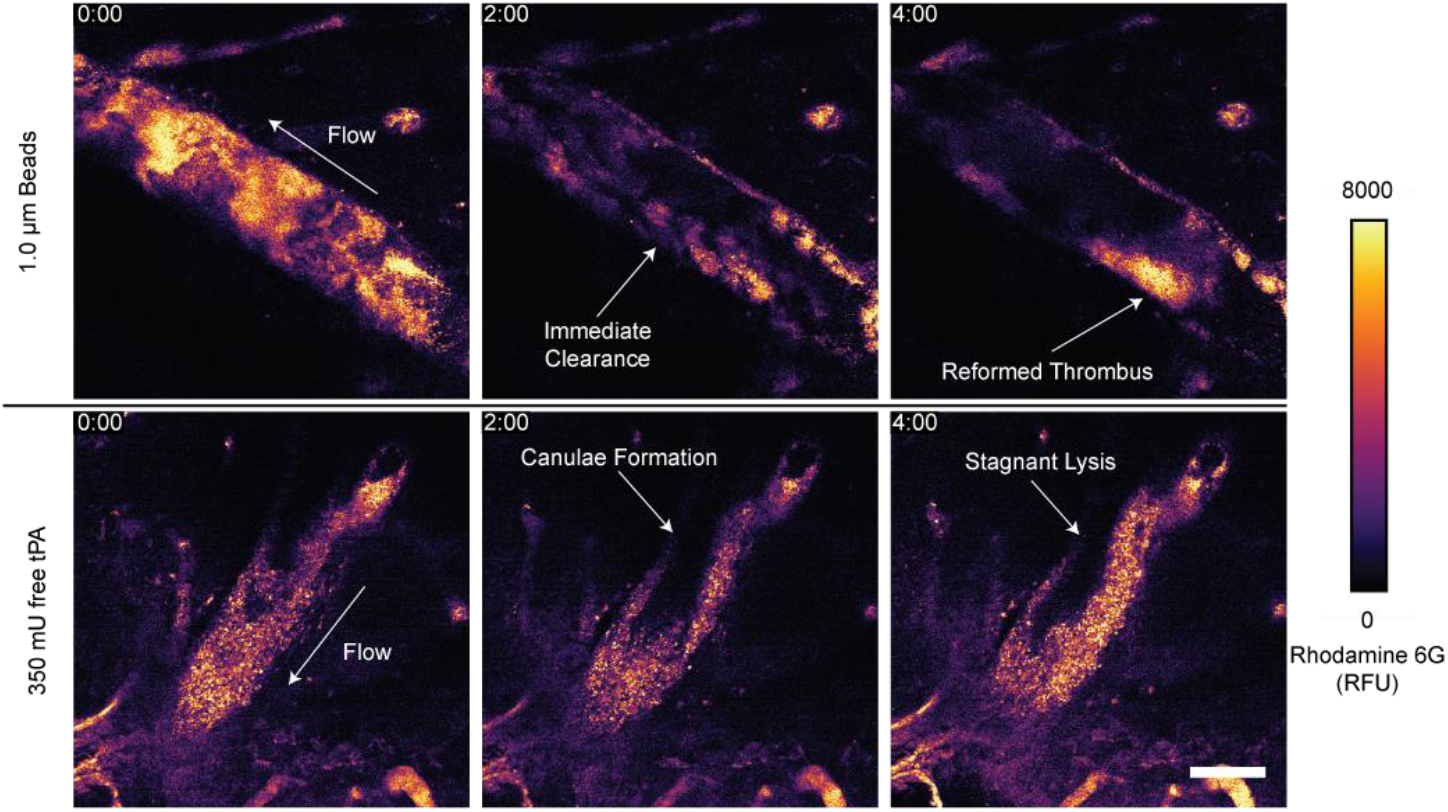
Thrombolysis observed by intravital two photon microscopy in the mouse middle cerebral artery. Blood cells were stained using rhodamine 6G (2.5 mg/kg) where the higher intensity portions of the heat mapped images indicate higher cell density within the thrombus. Treatments of 4.5 mU 1 μm tPA-beads and 350 mU free tPA were injected through the tail vein. The direction of flow is illustrated with arrows in the initial frames. Scale bar = 50 μm. Time stamp is minutes:seconds.

## Discussion

Our findings demonstrate that immobilizing tPA on micrometer-scale beads markedly enhances local plasmin generation compared to smaller beads or free tPA. The larger bead size is critical, as it supports a high local plasmin flux that overcomes transport limitations and facilitates a self-propagating wave of fibrinolysis. These same tPA-beads can rapidly re-establish blood flow in a murine model of ischemic stroke at a dose almost two orders-of-magnitude less than that required for free tPA. The immobilization of tPA to a bead is fundamentally different than the mechanism by which free tPA generates plasmin. In the case of free tPA, a complex between tPA-fibrin-plasminogen localizes plasmin generation to the fibrin fiber and minimizes its inhibition by antiplasmin. In the case of immobilized tPA, a similar complex can form with fibrinogen or fibrinogen degradation products to serve as a co-factor; however, the plasmin generated on the bead surface must be transported to fibrin fibers. The implications of these differences are discussed below.

By standardizing tPA activity across our experiments, we can directly compare the effects of bead size on plasmin generation. Our results show that 1 μm tPA-beads generate significantly more plasmin than both 0.1 μm beads and free tPA, highlighting the importance of bead size in enhancing fibrinolytic activity. An approximately 200-fold higher particle density of 0.1 μm beads is needed to match the tPA activity of 1 μm beads, which is twice as many as expected based on surface area. This suggests a difference in tPA enzymatic activity on the two sized beads, as supported by observations that, at equivalent tPA activities, 1 μm beads have higher plasmin generation than 0.1 μm beads, with a profound difference in tPA co-factors, fibrinogen and fibrin degradation products. Potential explanations for these observations include differences in curvature and steric limitations for substrate and co-factor binding. A reaction-diffusion model of plasmin generation on tPA-beads, even with comparable reaction kinetics between the two sizes, predicts that plasmin generation is reaction limited on the smaller beads in agreement with our experimental observations.

Comparing free tPA and 1 μm tPA-beads shows significantly faster and different lysis patterns *in vitro*. Plasmin generation with co-factors for a given tPA activity is slightly higher, 10-15%, for 1 μm tPA-beads compared to free tPA. Yet, the rate of internal fibrinolysis, as measured by turbidity, and external fibrinolysis, as measured by the ball sedimentation assay, are markedly faster for 1 μm tPA-beads. Examining fibrinolysis at the bead-scale suggests that this marked difference is due to a self-propagating fibrinolysis front that emanates away from the beads as a halo decorated with high concentration of plasmin(ogen). These halos grow linearly with time to the size for 100’s of micrometers, much faster than a diffusion mediated process. One potential explanation is the bundling of fibrin fibers upon lysis; fibrin fibers have an inherent tension and, when cut, retract like a taut rope forming bundles of fibers.(*40*–*43*) The thickening band of fluorescence in the growing halos that form around tPA-beads could be indicative of such fiber bundling. The proximity of fibers in a bundle could allow for plasmin to unbind and rebind between fibers at a favorable rate compared to diffusion or inhibition by antiplasmin or, alternatively enhance the rate of crawling between binding sites.(*16, 44*)

At a 77-fold lower dose, 1 μm tPA beads result in faster and more complete lysis than free tPA. This remarkable difference likely reflects the high plasmin generation on the bead surface that both overcomes antiplasmin inhibition and appears to bind to adjacent fibrin fibers at high concentrations *in vitro*. The thrombi formed in the photothrombotic model used in this study are porous enough to allow for interstitial plasma flow as evidenced by the entrainment of 1 μm particles through the thrombi. This could explain why *in vivo* lysis is much faster than *in vitro* lysis; the nonzero interstitial velocity through the thrombus replenishes plasminogen near the bead surface, which our calculations suggest is depleted in a purely diffusive environment. The thrombi were only 10 min old before treatment was introduced and as such, they were fibrin rich, porous, and potentially not highly contracted by platelets. Older clots that are more cellular and retracted will likely be harder to lyse; however, we have previously shown *in vitro* and in zebrafish models of thrombosis that tPA immobilized to superparamagnetic beads and assembled into wheel-like microbots can be directed to and driven into platelet-rich thrombi.(*45, 46*) The pattern of lysis was different between free tPA and tPA-beads. Free tPA initiates small channels that slowly widen at the blood-thrombus interface indicative of more permeable regions in the middle of the clot.(*21, 47, 48*) tPA-beads however lyse the clot from the inside-out resulting in failure-like events where the entire thrombus loses mechanical fidelity and dislodges from the vessel wall.

In summary, this study shows that immobilized tPA mediated plasmin generation at the micrometer scale supports rapid fibrinolysis *in vitro* and *in vivo*. It does so by overcoming mass transfer barriers, inhibition by antiplasmin, and populating fibrin fibers with a plasmin concentration that can sustain a self-propagating wave of fibrinolysis. As such the lysis patterns are fundamentally different than conventional tPA therapies where plasmin and tPA are co-localizing but also competing for binding sites on fibrin fibers. By achieving rapid recanalization at significantly lower tPA doses, this approach has the potential to minimize the risk of hemorrhagic complications associated with high-dose tPA therapy. The ability to efficiently dissolve clots with lower drug amounts could improve safety profiles and expand the therapeutic window for patients with acute ischemic stroke.

## Materials and Methods

### Materials

Dynabeads^®^ MyOne™ Carboxylic Acid (cat# 65011), and Fluospheres™ Carboxylate-modified microspheres 1.0 μm diameter (cat# F8823), 0.1 μm diameter (cat# F8803), and 96 well optical bottom plates (cat# 265301) were obtained from Thermo Fisher Scientific (Waltham, MA). 1-ethyl-3-[3-dimethylaminopropyl] carbodiimide hydrochloride (EDC) (cat# c1100-100mg) and N-hydroxysulfosuccinimide (Sulfo-NHS) (cat# c1102-1gm) were obtained from Proteochem (Hurricane, UT). Recombinant human tissue plasminogen activator, Cathflo^®^ Activase^®^ (tPA, alteplase, Genentech, San Francisco, CA) was obtained from the Hemophilia and Thrombosis Center pharmacy. Chromogenic substrate S-2288 (cat# S820852) was obtained from Diapharma (West Chester, OH). Purified human lys-plasmin (cat# IHUPLMLYS1MG) and plasmin chromogenic substrate (cat# IAFPLMCGSLY25MG) was obtained from Innovative Research (Novi, MI). Pooled normal plasma (cat# 0010) was obtained from George King Bio-Medical Inc. 8×40mm clear shell vials (cat# 50-978-400) were obtained from FisherScientific (Watham, MA). Bovine serum albumin (BSA) (cat# A2153-100G), Rose Bengal (cat# 330000), bovine thrombin (cat# T4648), 2-(N-Morpholino) ethanesulfonic acid hydrate (MES) (cat# m8250), HEPES (cat# H3375) and sodium dodecyl sulfate (SDS) (cat# 7910) was obtained from Sigma Aldrich (St. Louis, MO). Human fibrinogen depleted of plasminogen, von Willebrand Factor and fibronectin (FIB3), human glu-plasminogen (HPG 2001) and α2-antiplasmin (α2AP) was obtained from Enzyme Research Laboratories (South Bend, IN). Dade Innovin (tissue factor) was obtained from Siemens. HEPES buffered saline (HBS) was made with 20 mM HEPES and 100 mM sodium chloride pH 7.5. HEPES buffer used for purifying tPA was made with 0.3M HEPES at pH 7.4. Coupling buffer was made from 50 mM MES and 0.01% SDS at pH 6.5.

### Synthesis and characterization of tPA-beads

The lyophilized alteplase was dissolved in HEPES buffer (1mg/mL) and transferred to a slide-a-lyzer (MWCO 10,000) (Pierce, cat# 66455) dialysis cassette and dialyzed in 500 mL HEPES buffer at 4°C for 4 hr. This is necessary to remove arginine from the Cathflo^®^ formulation which can interfere with the linkage chemistry. After dialysis, the solution was removed from the cassette and the concentration measured using spectrophotometry (NanoDrop One, Thermofisher) and diluted to 1mg/mL in HEPES buffer. The solution was stored at -80 ºC. Both 0.1 and 1 μm beads were functionalized in 1 mg batches. Beads were added to microcentrifuge tubes and washed by suspending in coupling buffer (50 mM MES buffer, 0.01% SDS, pH 6.5) centrifuging at 3000g for 5 min and removing the supernatant. Next, EDC (100 μL, 40 mg/mL) and sulfo-NHS (100 μL, 40 mg/mL), both in coupling buffer, were added to the beads and mixed on a rotator for 15 min. The beads were then washed with coupling buffer to remove excess EDC/NHS. The dialyzed tPA (30 μg in 200 μL coupling buffer) was added to the bead suspension and gently inverted until the beads were fully dispersed. The beads were then mixed on a rotator at 4°C. After reacting for 4 hr, the beads were washed once with coupling buffer and blocked using 2% BSA in 0.9% saline for 5 min, washed three times to remove any excess tPA or BSA and stored in 0.9% saline. All beads were either used immediately or frozen at -80 °C. All steps outlined above were produced using sterile solutions in a sterile environment.

### tPA activity measurement with colorimetric substrate

Tris Buffer (50 μL, pH 8.4) and 50 μL of the free tPA (0-2 μg/mL final) or tPA-beads (diluted 100x, ∼10 μg beads/mL final) was added to each well of a 96 well optical bottom plate and mixed by pipetting up and down. Next, the tPA colorimetric substrate S-2288 (50 μL, 2.5 mM final, Km = 0.3-1 mM; kcat = 26-28 s-1) was added to each well and the absorbance was measured at 405 nm every min for 30 min. The activity defined the slope of the absorbance versus time curve (abs/min) over 20 min and multiplied by 313 to get the activity in units of U/L according to the manufacturer’s instructions.

### Fibrinolysis turbidity assay

Normal pooled plasma (50 μL), tissue factor (Dade Innovin, 50 μL, 1.25 pM final), and CaCl2 (50 μL, 8.3 mM final) and free or tPA (50 μL; 2.25, 45 U/mL) or tPA-beads (50 μL, 2.25 U/mL) in HEPES buffered saline (HBS) were added into a well of a 96 well plate and absorbance was measured at 450 nm at 37°C. Alternatively, to determine the effect of anti-plasmin presence, a measurement was made using a purified system of fibrinogen (40 μL, 1mg/mL final), glu-plasminogen (40 μL, 1 μM final), thrombin (40 μL, 2 U/mL final), anti-plasmin (40 μL, 1 μM final), and tPA or tPA-beads (40 μL).

### Formation of fibrin degradation products

Fibrin degradation products (FDP) were generated with fibrinogen (2 mg/mL), human alpha-thrombin (2 U/mL), and Lys-plasmin (1 μg/mL) for 16 hr at 37°C, followed by the addition of Pefabloc (0.05 mM final). The solution mixture was centrifuged at 6000g to remove any insoluble products.(*49*) The FDPs were diluted using HBS until the absorbance at 280 nm was 1.6, corresponding to a concentration of 1 mg/mL.

### Plasmin generation assay

Reaction mixtures containing human glu-plasminogen (1 μM), plasmin chromogenic substrate (0.2 mM) and tPA (2.25, 45 mU/mL) or tPA-beads (2.25 mU/mL; 0.1, 1 μm diameter) were placed in a 96 well assay plate. Fibrinogen (0.5 mg/mL) or FDPs (0.5 mg/mL) were added as co-factors in some wells. The absorbance at 405 nm was measured every minute for 3 hr.

### Confocal microscopy of single bead fibrin lysis

Fibrinogen (1 mg/mL final), AlexaFluor 647-tagged fibrinogen (40 μg/mL final), 0.1 μm and 1.0 μm tPA-beads (0.5 μg/mL final) or free tPA (0.4 nM) were mixed and followed by addition of thrombin (2 U/mL final), glu-plasminogen (0.9 μM), and Pacific Blue-tagged glu-plasminogen (0.1 μM) and placed on a cover slip. Fibrin fibers and fluorescent beads were observed using a confocal microscope (Olympus IX83 [60× objective, NA = 1.35, oil immersion]; Yokogawa CSU-W1 spinning disk confocal unit; Thorlabs laser lines—405 nm, 488 nm, and 640 nm; and Hamamatsu ORCA-Flash4.0 Digital CMOS camera) and recorded every 10 s until the fibrin network was lysed.

### Reaction-diffusion model of plasmin generation

The spatial-temporal evolution of plasmin formed on tPA beads was modeled as a three-component reaction-diffusion system including plasminogen, plasmin, and antiplasmin using three partial differential equations (see Supplemental Materials). These differential equations and their initial and boundary conditions were nondimensionalized and solved using the method of lines using custom Python scripts. Two Dahmköhler numbers dictate the concentration profile of plasmin; Da_bead_ which describes the rate of tPA mediated activation of plasminogen on the bead surface relative to the rate of plasminogen diffusion to the surface, and Da_antiplasmin_ which describes the rate of plasmin-antiplasmin binding compared to the rate of plasmin diffusion away from the surface.

### Ball Sedimentation Assay

Fibrinogen (2 mg/mL final), plasminogen (0.2 μM final) and thrombin (2 U/mL final) in HBS were added to an 8 mm diameter x 40 mm long vial and allowed to gel for 5 min at 37°C before a 2 mm steel bead was placed on the top of the clot. The lysis was initiated by addition of tPA or tPA-beads (100 μL) to the top surface of the fibrin gel for the following conditions: 4.5 mU tPA on 0.1 μm or 1.0 μm beads, or 4.5 mU of free tPA and HBS as a control. The vials were kept at 37°C and a time lapse image was recorded every min for 6 hr using a Canon EOS M200 camera. The time it took for the ball to fall to the bottom and the distance traveled over time were measured and reported.

### Murine photothrombotic stroke model

All procedures involving animals were reviewed and approved by the Institutional Animal Care and Use Committee of the University of Colorado (IACUC protocol number: 221). Before anesthesia, mice were warmed to dilate the tail vein and a temporary catheter was placed for later intravenous injections. Mice C57BL/6J mice (age 10-12 weeks) were deeply anesthetized using 3-5% isoflurane and maintained using 1-3% isoflurane. Mouse body temperature was maintained at 37°C. Animals were secured to a flat plate and visualized using a dissecting microscope (Leica M60). The left middle cerebral artery (MCA) was exposed using a process described previously.(*50, 51*) The left temporal muscle was transected to expose the skull. The proximal branch of the MCA was located and a 1.8 mm portion of the skull directly above the vessel was removed using a trephine (Fine science tools #18004-18). A bare fiber optic cable (200 μm diameter) connected to a 530 nm LED (Thorlabs M530F2) was positioned over the MCA vessel and a laser doppler flowmeter (LDF, MoorLabs DRT4) probe was positioned downstream to measure the relative blood flow. Blood flow measurements were collected for the entirety of the experiment. Each mouse was injected with Rose Bengal (40 mg/kg) and bovine thrombin (80 U/kg)(*39*). Immediately after injection, the MCA was illuminated at full power and the occlusion was confirmed by a drop in the flow measured by the LDF. The MCA was illuminated for 10 min and monitored to verify the stability of the clot. Treatment of the clots started ∼10 min after the beginning of the illumination. A bolus (∼120 μl) of saline, free tPA (4.5, 350 mU tPA), tPA-beads (4.5 mU tPA, diameter 0.1, 1 μm), or unfunctionalized beads (diameter 0.1, 1 μm) were delivered using the catheter and the LDF was monitored for 30 min to measure blood flow.

### Intravital microscopy of thrombolysis

The same procedures for the murine photothrombotic stroke model were used for intravital microscopy studies with some modification: To create a cranial window, the distal branch of the MCA was traced to the top of the head and a portion of the skull was removed using a trephine. An acrylic head plate with a viewing hole was centered and securely glued around the cranial window. The window was covered with sterile saline to prevent it from drying out. The mouse was then transported to the microscope and secured using a custom stereotaxic holder for the headplate. Mice were maintained at 37°C for the entire procedure. Isoflurane was supplemented with 50 mg/kg alpha-chloralose and 750 mg/kg urethane via peritoneal injection to provide prolonged anesthesia for the entirety of the imaging process. In addition to Rose Bengal and bovine thrombin, rhodamine 6g (2.5 mg/kg) was co-injected to visualize blood cells within the thrombus. The vessel was illuminated for 10 min using the 530 nm fiber optic LED to produce a thrombus and then the mouse was immediately positioned for viewing with a Bruker Ultima Investigator multiphoton microscope using Prairie View software and a Spectra-Physics Mai Tai® eHP DeepSee™ ultrafast laser.The system integrates two imaging channels with high sensitivity GaASP photomultiplier tubes.Excitation was done at 810 nm and green and red fluorescence emission were collected through 525/70-nm and 595/50-nm bandpass filters.The formed thrombus was located and recorded to ensure complete occlusion of the vessel. After formation of the thrombus, saline, free tPA, or tPA beads were injected at the same doses described above, and the lysis was recorded for 20 min. In some experiments, nonfunctionalized beads (0.1, 1 μm) labeled with FITC of the same size were added to visualize their distribution in and around the thrombus.

### Statistical Analysis

Results measured for significant differences were tested using the Shapiro-Wilk normality test. All data presented here was determined to have normal distribution. A two-way ANOVA was used to identify significant differences between the slopes of plasmin generation. Multiple comparisons were made between conditions using the same template or the same tPA source using Fisher’s least significant difference method.

## Author contributions

Conceptualization: KBN, DWMM

Methodology: MJO, KBN, DWMM

Validation: MJO, KBN, DWMM

Formal Analysis: MJO, KBN

Investigation: MJO, KBN

Resources: FD, NQ, DL, EJS, KBN, DWMM

Visualization: MJO, KBN, DWMM

Supervision: KBN, DWMM

Writing—original draft: MJO, KBN, DWMM

Writing—review & editing: MJO, KBN, DWMM, EJS, DL, FD, NQ

## Data and materials availability

All data are available in the main text or the supplementary materials.

