## Supplementary Materials for "Harnessing micrometer-scale tPA beads for high plasmin generation and accelerated fibrinolysis"

Matthew J. Osmond *et al.*

**This PDF file includes:**

Supplementary Text
Figs. S1 to S4

**Other Supplementary Materials for this manuscript include the following:**

Movies S1 to S3

### Supplementary Text

The derivation and numerical solution of the reaction-diffusion equations governing the conversion of plasminogen to plasmin catalyzed by tissue plasminogen activator (tPA) immobilized on spherical beads is described below. This model is used to aid in the interpretation of the single bead fibrinolysis assays. The model also includes the inhibition of plasmin by antiplasmin in the bulk, aiding in the interpretation of fibrinolysis experiments measured by turbidity with and without antiplasmin.

Starting from Michaelis-Menten kinetics, the equations are simplified under the assumption that the Michaelis constant ($K_m = 27\,\mu$M) is much greater than the plasminogen concentration (1-2 $\mu$M), leading to first-order kinetics at the bead surface (*6*). The equations are then nondimensionalized, introducing two Damköhler numbers ($\mathrm{Da}_{\mathrm{bead}}$ and $\mathrm{Da}_{\mathrm{ap}}$) to characterize the relative rates of reaction and diffusion.

We consider three species: plasminogen (Pgn), plasmin (Pln), and antiplasmin (AP). The concentration of each species depends on both time $t$ and radial position $r$ from the center of the bead. The concentration profiles of plasminogen, plasmin, and antiplasmin evolve according to the following equations:

$$\frac{\partial C_{\mathrm{Pgn}}}{\partial t} = D_{\mathrm{Pgn}} \left( \frac{1}{r^2} \frac{\partial}{\partial r} \left( r^2 \frac{\partial C_{\mathrm{Pgn}}}{\partial r} \right) \right) \tag{1}$$

$$\frac{\partial C_{\mathrm{Pln}}}{\partial t} = D_{\mathrm{Pln}} \left( \frac{1}{r^2} \frac{\partial}{\partial r} \left( r^2 \frac{\partial C_{\mathrm{Pln}}}{\partial r} \right) \right) - k_{\mathrm{ap}} C_{\mathrm{Pln}} C_{\mathrm{AP}} \tag{2}$$

$$\frac{\partial C_{\mathrm{AP}}}{\partial t} = D_{\mathrm{AP}} \left( \frac{1}{r^2} \frac{\partial}{\partial r} \left( r^2 \frac{\partial C_{\mathrm{AP}}}{\partial r} \right) \right) - k_{\mathrm{ap}} C_{\mathrm{Pln}} C_{\mathrm{AP}} \tag{3}$$

Here, $D$ represents the diffusion coefficients and $k_{\mathrm{ap}}$ is the rate constant for the reaction between plasmin and antiplasmin.

At time $t = 0$, the initial concentrations are:

$$C_{\mathrm{Pgn}}(r, 0) = C_{\mathrm{Pgn0}} \tag{4}$$

$$C_{\mathrm{Pln}}(r, 0) = 0 \tag{5}$$

$$C_{\mathrm{AP}}(r, 0) = C_{\mathrm{AP0}} \tag{6}$$

with boundary conditions at the bead surface ($r = R$):

$$-D_{\text{Pgn}} \left. \frac{\partial C_{\text{Pgn}}}{\partial r} \right|_{r=R} = -k_{\text{bead}} C_{\text{Pgn}}(R, t) \tag{7}$$

$$-D_{\text{Pln}} \left. \frac{\partial C_{\text{Pln}}}{\partial r} \right|_{r=R} = +k_{\text{bead}} C_{\text{Pgn}}(R, t) \tag{8}$$

$$-D_{\text{AP}} \left. \frac{\partial C_{\text{AP}}}{\partial r} \right|_{r=R} = 0 \tag{9}$$

At the outer boundary ($r = R_{\text{max}}$), the concentrations are maintained at bulk values:

$$C_{\text{Pgn}}(R_{\text{max}}, t) = C_{\text{Pgn0}} \tag{10}$$

$$C_{\text{Pln}}(R_{\text{max}}, t) = 0 \tag{11}$$

$$C_{\text{AP}}(R_{\text{max}}, t) = C_{\text{AP0}} \tag{12}$$

To simplify the equations and reduce the number of parameters, we introduce dimensionless variables:

$$r^* = \frac{r}{R} \tag{13}$$

$$t^* = \frac{t D_{\text{Pgn}}}{R^2} \tag{14}$$

$$C^*_{\text{Pgn}} = \frac{C_{\text{Pgn}}}{C_{\text{Pgn0}}} \tag{15}$$

$$C^*_{\text{Pln}} = \frac{C_{\text{Pln}}}{C_{\text{Pgn0}}} \tag{16}$$

$$C^*_{\text{AP}} = \frac{C_{\text{AP}}}{C_{\text{AP0}}} \tag{17}$$

with dimensionless parameters defined as:

$$\alpha_{\text{Pln}} = \frac{D_{\text{Pln}}}{D_{\text{Pgn}}} \tag{18}$$

$$\alpha_{\text{AP}} = \frac{D_{\text{AP}}}{D_{\text{Pgn}}} \tag{19}$$

The Damköhler numbers, which characterize the ratio of reaction rate to diffusion rate, are introduced as:

$$\mathrm{Da}_{\mathrm{bead}} = \frac{k_{\mathrm{bead}} R}{D_{\mathrm{Pgn}}} \tag{20}$$

$$\mathrm{Da}_{\mathrm{ap}} = \frac{k_{\mathrm{ap}} C_{\mathrm{Pgn0}} R^2}{D_{\mathrm{Pgn}}} \tag{21}$$

Using these substitutions, the dimensionless governing equations become:

$$\frac{\partial C^*_{\mathrm{Pgn}}}{\partial t^*} = \frac{1}{(r^*)^2} \frac{\partial}{\partial r^*} \left( (r^*)^2 \frac{\partial C^*_{\mathrm{Pgn}}}{\partial r^*} \right) \tag{22}$$

$$\frac{\partial C^*_{\mathrm{Pln}}}{\partial t^*} = \alpha_{\mathrm{Pln}} \left[ \frac{1}{(r^*)^2} \frac{\partial}{\partial r^*} \left( (r^*)^2 \frac{\partial C^*_{\mathrm{Pln}}}{\partial r^*} \right) \right] - \mathrm{Da}_{\mathrm{ap}} C^*_{\mathrm{Pln}} C^*_{\mathrm{AP}} \tag{23}$$

$$\frac{\partial C^*_{\mathrm{AP}}}{\partial t^*} = \alpha_{\mathrm{AP}} \left[ \frac{1}{(r^*)^2} \frac{\partial}{\partial r^*} \left( (r^*)^2 \frac{\partial C^*_{\mathrm{AP}}}{\partial r^*} \right) \right] - \mathrm{Da}_{\mathrm{ap}} C^*_{\mathrm{Pln}} C^*_{\mathrm{AP}} \tag{24}$$

with dimensionless Initial Conditions:

$$C^*_{\mathrm{Pgn}}(r^*, 0) = 1 \tag{25}$$

$$C^*_{\mathrm{Pln}}(r^*, 0) = 0 \tag{26}$$

$$C^*_{\mathrm{AP}}(r^*, 0) = 1 \tag{27}$$

and boundary conditions at $r^* = 1$:

$$- \left. \frac{\partial C^*_{\mathrm{Pgn}}}{\partial r^*} \right|_{r^*=1} = \mathrm{Da}_{\mathrm{bead}} C^*_{\mathrm{Pgn}}(1, t^*) \tag{28}$$

$$-\alpha_{\mathrm{Pln}} \left. \frac{\partial C^*_{\mathrm{Pln}}}{\partial r^*} \right|_{r^*=1} = -\mathrm{Da}_{\mathrm{bead}} C^*_{\mathrm{Pgn}}(1, t^*) \tag{29}$$

$$-\alpha_{\mathrm{AP}} \left. \frac{\partial C^*_{\mathrm{AP}}}{\partial r^*} \right|_{r^*=1} = 0 \tag{30}$$

and at $r^* = r^*_{\mathrm{max}}$:

$$C^*_{\mathrm{Pgn}}(r^*_{\max}, t^*) = 1 \tag{31}$$

$$C^*_{\mathrm{Pln}}(r^*_{\max}, t^*) = 0 \tag{32}$$

$$C^*_{\mathrm{AP}}(r^*_{\max}, t^*) = 1 \tag{33}$$

With the nondimensional equations established, we calculate the Damköhler numbers to characterize the relative importance of reaction and diffusion. We begin by estimating the necessary parameters.

Assuming full monolayer coverage of tPA on the bead:

$$A_{\mathrm{tPA}} = \pi \left(\frac{d_{\mathrm{tPA}}}{2}\right)^2 \approx \pi(2.5 \times 10^{-9}\,\mathrm{m})^2 \approx 1.96 \times 10^{-17}\,\mathrm{m}^2 \tag{34}$$

$$[E]_{\mathrm{surface}} = \frac{1}{A_{\mathrm{tPA}} N_A} \approx \frac{1}{1.96 \times 10^{-17} \times 6.022 \times 10^{23}} \approx 8.48 \times 10^{-8}\,\mathrm{mol/m}^2 \tag{35}$$

and then we can calculate $V_{\max}$ and $k_{\mathrm{bead}}$ using $k_{\mathrm{cat}} = 0.3\,\mathrm{s}^{-1}$ *6*:

$$V_{\max} = k_{\mathrm{cat}}[E]_{\mathrm{surface}} = (0.3)(8.48 \times 10^{-8}) = 2.544 \times 10^{-8}\,\mathrm{mol/(m^2 \cdot s)} \tag{36}$$

$$k_{\mathrm{bead}} = \frac{V_{\max}}{K_m} = \frac{2.544 \times 10^{-8}}{27 \times 10^{-6}} = 9.422 \times 10^{-4}\,\mathrm{m/s} \tag{37}$$

We use the Stokes-Einstein equation to estimate diffusion coefficients at $T = 37°\mathrm{C}$ ($\eta = 0.691 \times 10^{-3}\,\mathrm{Pa \cdot s}$):

$$D_{\mathrm{Pgn}} \approx 1.106 \times 10^{-10}\,\mathrm{m}^2/\mathrm{s} \tag{38}$$

$$D_{\mathrm{Pln}} \approx 1.128 \times 10^{-10}\,\mathrm{m}^2/\mathrm{s} \tag{39}$$

$$D_{\mathrm{AP}} \approx 1.208 \times 10^{-10}\,\mathrm{m}^2/\mathrm{s} \tag{40}$$

For simplicity, we use $D_{\mathrm{Pgn}} = D_{\mathrm{Pln}} = D_{\mathrm{AP}} = 1.15 \times 10^{-10}\,\mathrm{m}^2/\mathrm{s}$.

We can now calculate the Damköhler number at $T = 37°\mathrm{C}$ and for a bead radius $R = 0.05\,\mu\mathrm{m}$:

$$\mathrm{Da}_{\mathrm{bead}} = \frac{k_{\mathrm{bead}} R}{D_{\mathrm{Pgn}}} = \frac{(9.422 \times 10^{-4})(0.05 \times 10^{-6})}{1.15 \times 10^{-10}} \approx 0.4097 \tag{41}$$

$$\mathrm{Da}_{\mathrm{ap}} = \frac{k_{\mathrm{ap}} C_{\mathrm{Pgn0}} R^2}{D_{\mathrm{Pgn}}} = \frac{(2 \times 10^{7})(1 \times 10^{-3})(0.05 \times 10^{-6})^2}{1.15 \times 10^{-10}} \approx 0.4348 \tag{42}$$

For bead radius $R = 0.5\,\mu$m:

$$\mathrm{Da}_{\mathrm{bead}} \approx 4.097 \tag{43}$$

$$\mathrm{Da}_{\mathrm{ap}} \approx 43.478 \tag{44}$$

At $T = 20°$C, we adjust the rate constants using the Arrhenius equation and diffusion coefficients using the Stokes-Einstein equation. For $R = 0.05\,\mu$m:

$$\mathrm{Da}_{\mathrm{bead}} \approx 0.204 \tag{45}$$

$$\mathrm{Da}_{\mathrm{ap}} \approx 0.217 \tag{46}$$

For $R = 0.5\,\mu$m:

$$\mathrm{Da}_{\mathrm{bead}} \approx 2.04 \tag{47}$$

$$\mathrm{Da}_{\mathrm{ap}} \approx 21.7 \tag{48}$$

Table 1: Parameters used in the model

| Parameter | Description | Value | Units | Source/Derivation |
|---|---|---|---|---|
| $D_{\text{Pgn}}$ | Diffusion coefficient of plasminogen | $1.15 \times 10^{-10}$ | $m^2/s$ | Stokes-Einstein equation |
| $D_{\text{Pln}}$ | Diffusion coefficient of plasmin | $1.15 \times 10^{-10}$ | $m^2/s$ | Stokes-Einstein equation |
| $D_{\text{AP}}$ | Diffusion coefficient of antiplasmin | $1.15 \times 10^{-10}$ | $m^2/s$ | Stokes-Einstein equation |
| $V_{\text{max}}$ | Max surface reaction rate | $2.544 \times 10^{-8}$ | $\text{mol}/(\text{m}^2{\cdot}\text{s})$ | Calculated from $k_{\text{cat}}$ and $[E]_{\text{surface}}$ |
| $K_m$ | Michaelis-Menten constant | $27 \times 10^{-6}$ | $\text{mol/m}^3$ | Reference *6* |
| $k_{\text{bead}}$ | First-order rate constant | $9.422 \times 10^{-4}$ | m/s | $k_{\text{bead}} = V_{\text{max}}/K_m$ |
| $k_{\text{ap}}$ | Homogeneous reaction rate constant | $2 \times 10^7$ | $\text{m}^3/\text{mol}{\cdot}\text{s}$ | Reference *24* |
| $R$ | Radius of the bead | $0.05\,\mu\text{m}$, $0.5\,\mu\text{m}$ | m | Experimental |
| $C_{\text{Pgn0}}$ | Bulk concentration of plasminogen | $1 \times 10^{-3}$ | $\text{mol/m}^3$ | Experimental |
| $C_{\text{AP0}}$ | Bulk concentration of antiplasmin | $1 \times 10^{-3}$ | $\text{mol/m}^3$ | Experimental |

To solve the system of reaction-diffusion equations, we employ the method of lines (MOL), which discretizes the spatial domain while leaving time continuous, transforming the partial differential equations into ordinary differential equations (ODEs). The spatial domain from $r^* = 1$ to $r^* = r^*_{\text{max}}$ is discretized into $N$ nodes using a finite difference grid. Central difference schemes are used for the spatial derivatives, converting the PDEs into a system of ODEs with respect to time. The Laplacian operator in spherical coordinates is discretized as:

$$\left(\frac{1}{r^2}\frac{\partial}{\partial r}\left(r^2\frac{\partial C}{\partial r}\right)\right)_i = \frac{1}{r_i^2}\left(r^2_{i+\frac{1}{2}}\frac{C_{i+1}-C_i}{\Delta r} - r^2_{i-\frac{1}{2}}\frac{C_i - C_{i-1}}{\Delta r}\right)\frac{1}{\Delta r} \tag{49}$$

where $r_i$ is the radial position at node $i$, and $\Delta r$ is the spatial step size.

The dimensionless initial conditions are applied directly to the initial state vector in the ODE solver:

$$C^*_{\mathrm{Pgn}}(r^*, 0) = 1 \tag{50}$$

$$C^*_{\mathrm{Pln}}(r^*, 0) = 0 \tag{51}$$

$$C^*_{\mathrm{AP}}(r^*, 0) = 1 \tag{52}$$

At the inner boundary ($r^* = 1$), the boundary conditions involve flux terms due to reactions. We implement these using one-sided finite differences. For example, the boundary condition for plasminogen becomes:

$$\frac{C^*_{\mathrm{Pgn},2} - C^*_{\mathrm{Pgn},1}}{\Delta r^*} = -\mathrm{Da}_{\mathrm{bead}} C^*_{\mathrm{Pgn},1} \tag{53}$$

This equation can be rearranged to solve for $C^*_{\mathrm{Pgn},1}$ in terms of $C^*_{\mathrm{Pgn},2}$ and known parameters, ensuring that the boundary condition is satisfied. At the outer boundary ($r^* = r^*_{\mathrm{max}}$), we apply Dirichlet boundary conditions by setting:

$$C^*_{\mathrm{Pgn},N} = 1 \tag{54}$$

$$C^*_{\mathrm{Pln},N} = 0 \tag{55}$$

$$C^*_{\mathrm{AP},N} = 1 \tag{56}$$

The system of ODEs is integrated over time using the `solve_ivp` function from the `scipy.integrate` module in Python, with the method set to 'BDF' (Backward Differentiation Formula), suitable for stiff systems. At each time step, the boundary conditions are enforced, and the reaction terms are computed based on the current concentrations. Stiffness arising from rapid reactions and diffusion is managed by using implicit time integration methods ('BDF') and appropriate time step controls in the solver.

To determine when the system has reached steady state during the numerical solution of the reaction-diffusion equations, we employ a time derivative and concentration change criteria. The system is considered at steady state when the absolute value of the time derivatives of the dimensionless concentrations for all species $i$ at all spatial points $r^*$ falls below a small threshold $\epsilon$:

$$\left| \frac{\partial C_i^*(r^*, t^*)}{\partial t^*} \right| < \epsilon = 1 \times 10^{-6}$$

Steady state is achieved when the maximum change in dimensionless concentrations between successive time steps for all species $i$ across the spatial domain is less than a specified tolerance $\delta$:

$$\max_{r^*} \left| C_i^*(r^*, t^* + \Delta t^*) - C_i^*(r^*, t^*) \right| < \delta = 1 \times 10^{-6}$$

The time derivative criterion ensures that the rate of change of concentrations with respect to time is negligible throughout the spatial domain, indicating that transient behaviors have decayed. The concentration change criterion monitors the change in concentrations over a time step $\Delta t^*$. If the maximum change is below the tolerance $\delta$, it signifies that concentrations have stabilized.

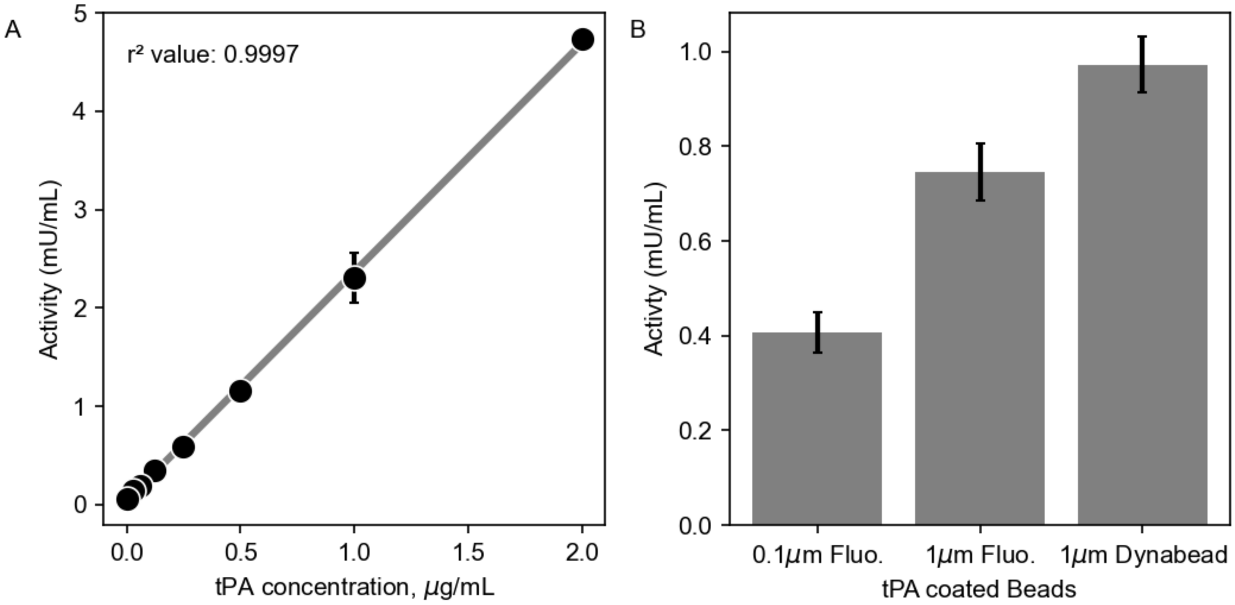


**Figure S1. Measured tPA activities**. (**A**). Measured activity of free tPA using chromogenic substrate S-2288 at concentrations from 0 to 2 μg/mL to generate a standard curve for comparison of tPA activity. (**B**). Relative activity of tPA functionalized beads compared to the activity of free tPA. All beads were diluted by 100-fold to reduce light scatter. The final bead densities for the 0.1μm bead concentration was $3.55 \times 10^{10}$ beads/mL; 1.0 μm bead concentration was $1.69 \times 10^{8}$ beads/mL. * $p<0.05$

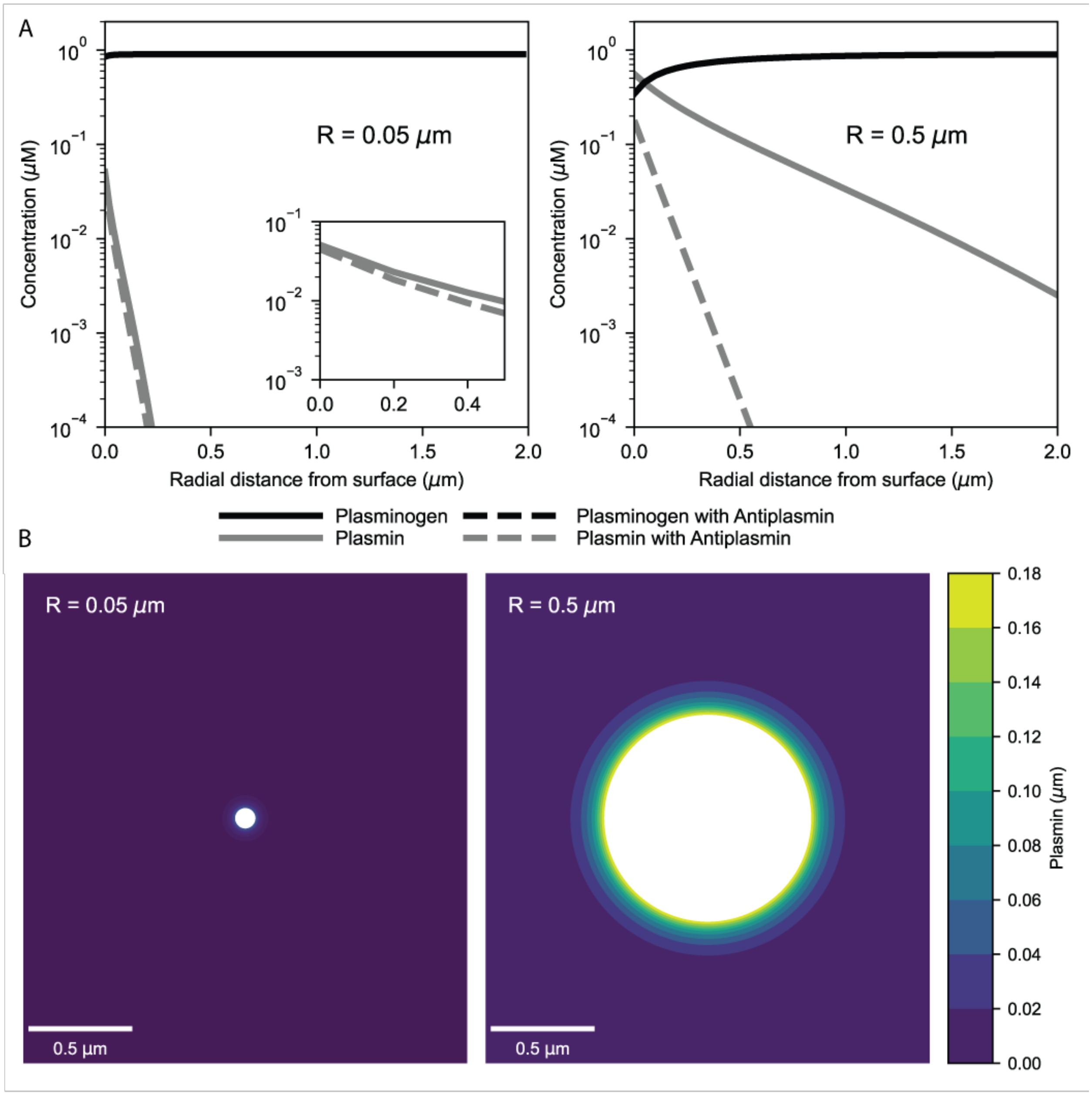


**Figure S2. Simulation of plasmin generation on beads with anti-plasmin.** A reaction-diffusion model of plasmin generation on tPA-beads was used to calculate the concentration profiles (**A**) of plasminogen and plasmin as a function of radial distance from the bead surface in the presence (1 μM) and absence of anti-plasmin and contour plots (**B**) of plasmin concentration on 0.1 and 1.0 μm tPA-beads in the presence of anti-plasmin. Reaction constants used were calculated for a temperature of 37°C. Initial concentrations of plasminogen and anti-plasmin were 0.9 μM and 1 μM respectively.

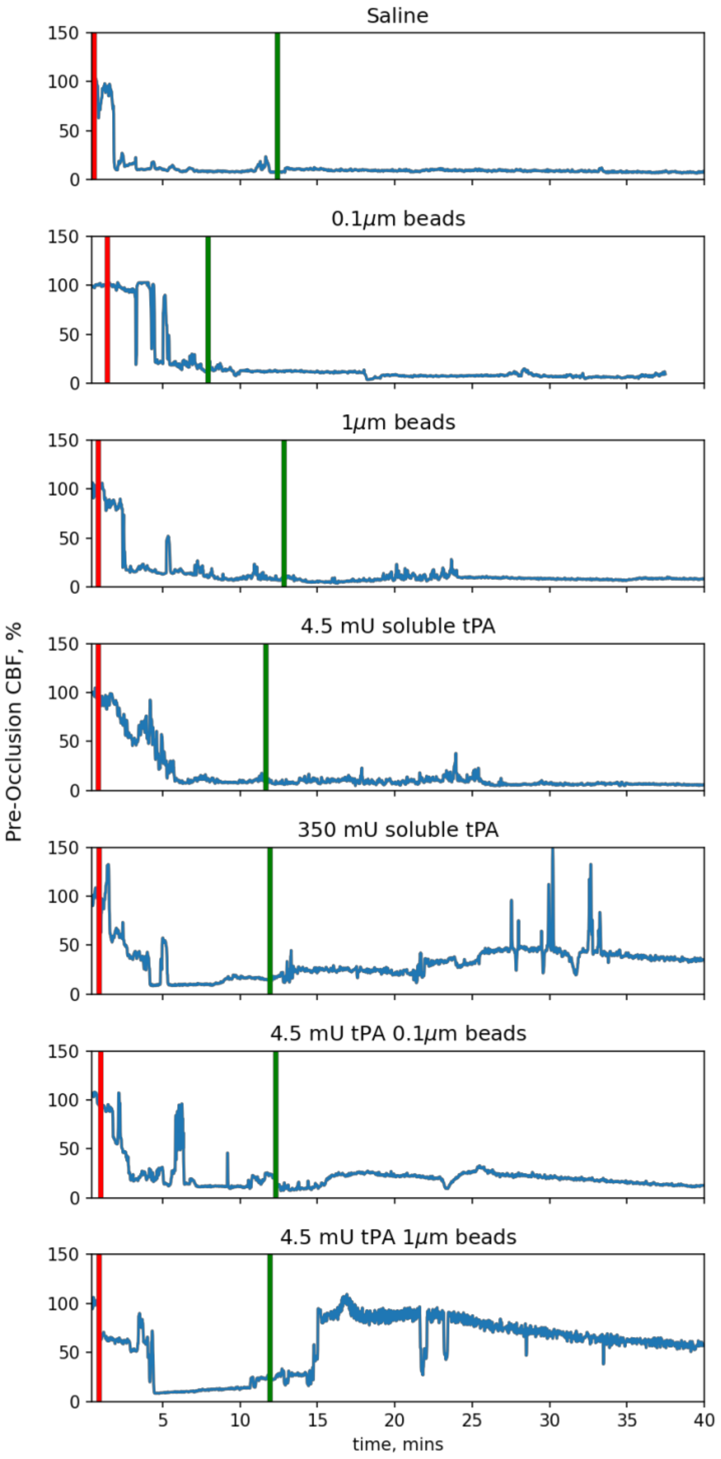


**Figure S3. Laser Doppler flowmeter measurements.** Representative laser doppler flowmeter measurements to illustrate the median condition for each of the above treatment conditions. Saline only control, 0.1 µm beads without tPA, 1.0 µm beads without tPA, 4.5 mU free tPA dose, 350 mU free tPA dose, 0.1 µm beads with 4.5 mU tPA dose and the 1.0 µm beads with 4.5 mU tPA dose.

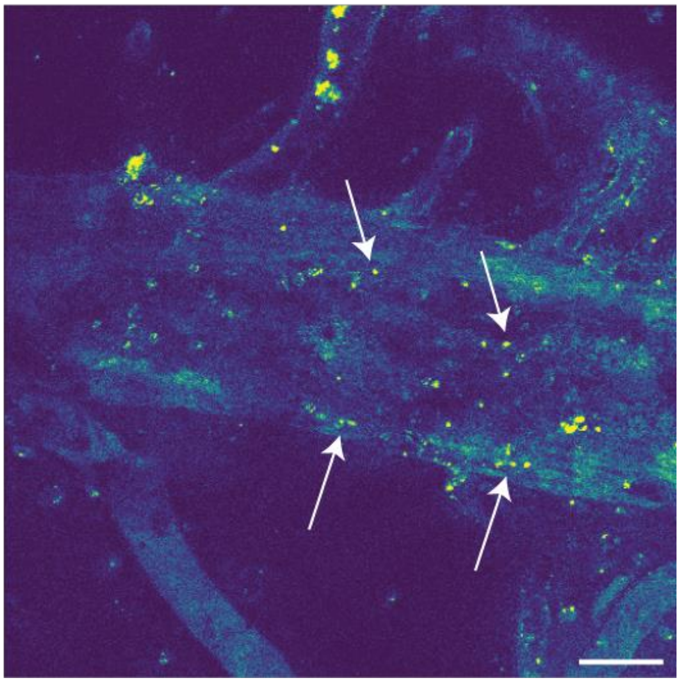

**Figure S4. Two-photon microscopy of occlusion.** Two photon micrograph of occluded mouse blood vessel with 1 μm fluorescent beads entrained. Scale bar = 50 μm.

**Movie S1. Growth of fibrinolytic halo around tPA-beads.** Confocal microscopy of the lysis front around a 1.0 μm tPA-bead. The fibrin network was labeled using 40μg/mL Alexa Fluor 647 labeled fibrinogen (1:25 labeled:unlabeled). Scale bar = 20 μm. Time stamp is hours:minutes:seconds.

**Movie S2. Incomplete recanalization of the mouse middle cerebral artery using free tPA.** Blood cells were stained using rhodamine 6g (2.5 mg/kg) where the higher intensity portions of the heat mapped images indicate higher cell density within the thrombus. A bolus of 350 mU free tPA was injected through the tail vein. Formation of canulae is visible through the center of the thrombus. Full recanalization never occurs. Scale bar = 50 μm. Time stamp is minutes:seconds.

**Movie S3. Spontaneous recanalization of the mouse middle cerebral artery using 1.0 μm beads.** Blood cells were stained using rhodamine 6g (2.5 mg/kg) where the higher intensity portions of the heat mapped images indicate higher cell density within the thrombus. A bolus of 1.0 μm beads with 4.5 mU of attached tPA was injected through the tail vein. Initially, some beads collected at the bottom right are visible. As time goes on, a spontaneous generation of plasmin is apparent which causes an immediate sloughing of the entire thrombus. Scale bar = 50 μm. Time stamp is minutes:seconds.
